# Directional motion sensitivity in people with Visual Snow Syndrome is modulated by the presence of trailing-type palinopsia

**DOI:** 10.1101/2024.09.25.614880

**Authors:** T.S. Obukhova, T.A. Stroganova, A.R. Artemenko, A.V. Petrokovskaia, E.V. Orekhova

**Affiliations:** Center for Neurocognitive Research (MEG Center), Moscow State University of Psychology and Education, Moscow, Russia; Sechenov First Moscow State Medical University of the Ministry of Health of the Russian Federation (Sechenov University), Moscow, Russia; Loginov Moscow Clinical Scientific Center, Moscow, Russia

**Keywords:** Visual Snow Syndrome (VSS), palinopsia, visual motion, spatial suppression

## Abstract

Visual Snow Syndrome (VSS) is characterized by visual perceptual distortions, potentially linked to increased neural excitability. We hypothesized that this hyperexcitability might affect motion direction sensitivity in VSS, particularly in those with trailing-type palinopsia (TTP), an atypical perception of visual motion.

Using a spatial suppression paradigm, we assessed motion duration discrimination thresholds for small (1°), medium (2.5°), and large (12°) high-contrast gratings in 23 VSS and 27 control participants. Spatial Suppression Index (SSI) quantified size-dependent increases in duration thresholds. Visual Discomfort Questionnaire scores and VSS symptom ratings including TTP, afterimages, photophobia, etc. were also collected.

VSS patients reported higher visual discomfort and perceptual disturbances. However, no group differences were found in duration thresholds or SSI. Notably, higher TTP scores were associated with lower duration thresholds, indicating a facilitatory effect of TTP on motion sensitivity.

These findings suggest that ‘visual snow,’ the core symptom of VSS, is not linked to abnormal directional sensitivity or center-surround suppression associated with visual motion. However, the dependence of directional sensitivity on TTP emphasizes the heterogeneity of VSS, which should be considered in future neurophysiological and clinical models.

## Introduction

Visual Snow Syndrome (VSS) is a neurological disorder characterized by the persistent small flickering dots across the entire visual field that resemble tiny snowflakes or noise in a poorly tuned analogue TV. Apart from the ‘snow’ people with VSS usually have a wide range of other visual and non-visual symptoms. The visual disturbances include photophobia, nyctalopia (night-blindness), palinopsia, enhanced entoptic phenomena, etc. (Puledda, Schankin, Digre, & Goadsby, 2018; Puledda, Schankin, & Goadsby, 2020). Among non-visual symptoms most common are tinnitus, migraine, impaired concentration, fatigue, and anxiety (Solly, Clough, Foletta, White, & Fielding, 2021; White, Clough, McKendrick, & Fielding, 2018).

It has been suggested that VSS is associated with neuronal hyperexcitability, which leads to perception of visual stimuli that do not exist (Bou Ghannam & Pelak, 2017; Lauschke, Plant, & Fraser, 2016). Evidence for hyperexcitability comes from several sources. For example, people with VSS experience strong visual discomfort when viewing bright light or high-contrast visual patterns (Brooks et al., 2022), a phenomenon that has been attributed to increased excitability in the visual pathways (Bargary, Furlan, Raynham, Barbur, & Smith, 2015). The reduced phosphene threshold during TMS stimulation of the occipital cortex found in individuals with VSS (Yildiz, Turkyilmaz, & Unal-Cevik, 2019) is also indicative of increased excitability. Other evidence pointing in the same direction includes shortened prosaccade latencies (Solly et al., 2020), lack of habituation of the visual evoked potentials (Luna, Lai, & Harris, 2018; Unal-Cevik & Yildiz, 2015; Yildiz et al., 2019), and increased metabolism in the lingual gyrus (Van Laere, Ceccarini, Gebruers, Goffin, & Boon, 2022). In addition, VSS is often comorbid with migraine and palinopsia, which are thought to be associated with altered excitation/inhibition (E/I) balance in the visual cortex (Aurora & Wilkinson, 2007; Gersztenkorn & Lee, 2015; Kalita, Uniyal, & Bhoi, 2016; Vecchia & Pietrobon, 2012). Some authors have suggested that neural excitability in VSS is the result of deficient inhibitory processes (Luna et al., 2018). However, there is no direct evidence of impaired inhibitory processes in VSS. In a TMS study, normal magnetic suppression of perceptual acuity was found, indicating normal inhibition in the visual cortex (Eren et al., 2021).

E/I imbalance in the visual cortex in patients with VSS may affect visual perception through altering center-surround antagonism, a phenomenon central to the ability to separate figure and/or motion from visual background (Tadin, Lappin, Gilroy, & Blake, 2003; Tadin et al., 2019). McKendrick et al (McKendrick et al., 2017) studied center-surround interactions in patients with VSS using center-surround contrast matching paradigm. Participants were presented with small moving gratings with and without an iso-orientated annular surround and compared the perceived contrast to an original reference. Participants with VSS showed less contrast suppression in the presence of the surround compared to controls. This effect may be due to weakened lateral inhibition, reduced feedback from extrastriate areas, or atypical feedforward connections from the lateral geniculate nucleus of the thalamus.

Another approach that is often used to study altered center-surround interactions in different patient and age groups is the spatial suppression paradigm initially proposed by Tadin et al. (Tadin et al., 2003). Although this paradigm has not been applied in VSS, it may shed light on the nature of E/I imbalance in this neurological condition. This approach exploits the observation that the duration of stimulation required to determine the direction of motion of high-contrast stimulus increases with stimulus size, reflecting the greater involvement of center-surround inhibition.

While discriminating of the direction of small-size objects (∼1° or less) is mainly influenced by E/I interactions within the primary visual cortex (area V1) (Angelucci et al., 2017; Shushruth et al., 2013) and/or MT (Schallmo et al., 2019), the paradoxically reduced ability to determine the direction of large (>1 degree) stimuli depends on top-down modulation of V1 inhibitory circuitry from the extrastriate areas (Angelucci et al., 2017; Nassi, Lomber, & Born, 2013). The mechanisms of this modulation are still debated (Liu, Miller, & Pack, 2018; Ozeki, Finn, Schaffer, Miller, & Ferster, 2009), but it effectively results in functional inhibition (reduced activity) in visual cortical areas (Schallmo et al., 2020).

Abnormalities in duration thresholds for the small and large stimuli may reflect alterations in different cortical mechanisms. *Elevated* duration thresholds for small high-contrast gratings have been observed in children with autism, which is compatible with an increase in receptive field sizes due to deficient local inhibition in V1 (Orekhova et al., 2023; Schauder, Park, Tadin, & Bennetto, 2017). On the other hand, patients with major depression disorder during remission had reduced duration thresholds for large-size gratings, presumably reflecting a deficit in top-down inhibitory mechanisms (Golomb et al., 2009).

In patients with Alzheimer’s Disease, thresholds are abnormally increased for small gratings and abnormally decreased for large gratings, suggesting deficits in both local and top-down inhibition (Zhuang et al., 2017). Thus, changes in duration thresholds for small or large stimuli (or both) may, albeit indirectly, indicate the level of the visual hierarchy at which E/I balance is disrupted in individuals with VSS.

In this study, we used the spatial suppression paradigm to investigate center-surround interactions in VSS. By investigating duration thresholds for the stimuli of different sizes we hoped to estimate different aspects of the E/I balance regulation in the visual cortex in patients with VSS.

Our second aim was to differentiate the effects of the visual snow (VS) on duration thresholds from the effect of palinopsia. Palinopsia is marked by the persistence of visual images after a stimulus has disappeared, manifesting as the retention of images from stationary or moving objects. In contrast to physiologically normal ‘negative’ afterimages, the images in palinopsia are almost always ‘positive’, i.e., isochromatic to the original stimulus (Gersztenkorn & Lee, 2015). According to Puledda et al. (2020), approximately 80% of patients with Visual Snow Syndrome (VSS) experience palinopsia. Although VSS often co-occurs with migraines, this prevalence is significantly higher than in migraine patients without VSS, where palinopsia occurs in about 10% of cases (Belcastro et al., 2011). Palinopsia has also been linked to increased cortical excitability (Gersztenkorn & Lee, 2015; Kalita et al., 2016), but the extent to which neural alterations in palinopsia overlap with and influence VSS remains unclear. We expected that the spatial suppression paradigm focused on motion perception might be particularly informative regarding E/I imbalance in VSS patients experiencing trailing-type palinopsia (TTP), a condition in which a moving object leaves behind a trail lasting several seconds.

## Methods

### Participants

Twenty-three patients recruited from an online community for people with VSS were included in the study. All patients were examined by a neurologist who collected information about their neurologic and visual symptoms. In 12 subjects the VS was present from an early age (∼ <10 years). In 11 subjects, it appeared after the age of 14 years. Six of these 11 participants attributed the onset of VS to stress. Subjects who acquired VS as a result of substance abuse were excluded from the study. All but two VSS participants were medication-free. Two participants were taking antidepressants in low doses (venlafaxin, 75 mg per day, or escitalopram, 5 mg per day, both longer than 3 months).

We also recruited 27 sex- and age-matched neurologically healthy control participants from the community. The characteristics of the subjects are presented in Table 1. All subjects were asked to complete the Russian version of the Visual Discomfort Scale (Conlon, Lovegrove, Chekaluk, & Pattison, 1999), which assesses unpleasant somatic and perceptual side effects of viewing patterns. Participants were also asked to complete the Russian version State-Trait Anxiety Inventory (Spielberger, 1983).

The local ethics committee approved the study and all subjects gave written informed consent.

**Table 1.** Characteristics of the participants.

|  | *VSS* | *Control* | *t, p* |
| --- | --- | --- | --- |
| N | 23 (12 males) | 27 (16 males) |  |
| Age | 26.1 +/- 5.4 | 26.9 +/- 4.2 | n.s. |
| Migraine | 17 (74%) | 0% |  |
| Migraine with aura | 6 (26%) | 0% |  |
| Tinnitus | 13 (57%) | 0% |  |
| Anxiety score: STAI (all life)* | 52.5 +/- 9.4 | 40.1 +/- 9.6 | t(48) = 4.57, p=0.00004 |
| Visual Discomfort score** | 26.2 +/- 13.0 | 8.5 +/- 7.9 | t(47) = 6.2 p<0.00001 |

\* Information was not available for one control participant.
\*\* Information was not available for one VS and one control participant.

Patients with VSS rated a range of their visual symptoms on a 5-point Likert scale (1 – absence of the symptom, 2 – very rare, 3 – rare, 4 – often, 5 – all the time). Table 2 summarizes the number of subjects who marked ‘4’ or ‘5’ on this scale for the symptoms most frequently associated with VSS. The presence of some of these symptoms was a necessary criterion for the diagnosis of VSS (Puledda et al., 2018).

**Table 2.** Visual phenomena and neurological symptoms in patients with VSS patients.

| *Visual symptoms* | *VSS* |
| --- | --- |
| VS from birth or early age (<10 years) | 10 (43%) |
| Palinopsia (I): afterimages | 13 (57%) |
| Palinopsia (II): trailing of moving objects | 8 (34%) |
| Enhanced entopic phenomena | 17 (74%) |
| Photophobia | 17 (74%) |
| Impaired night vision (Nyctalopia) | 14 (61%) |

The ‘object trailing’ visual symptoms were of the particular interest because of their potential relation to motion sensitivity assessed in our experimental paradigm. We therefore categorized palinopsia into two types: afterimages of static objects (type I) and trailing of moving objects (type II). All but one of the eight patients who scored 4 or 5 on the type-II palinopsia also scored 4 or 5 on the type-I palinopsia, but 6 subjects who scored 4 or 5 on the type-I palinopsia (afterimages) scored lower than 4 on the type-II palinopsia (trailing). Among patients with type-II palinopsia, three had migraine with aura, 2 had migraine without aura, and three had no migraine.

The majority of VSS subjects also had other visual symptoms, such as flashes of light (n=7, 30%), swirls or clouds with eyes closed (n=14, 61%), or other forms of unusual visual phenomena (n=13, 57%).

### Stimuli and Paradigm

Visual stimuli were presented using PsychoToolbox (Brainard). We used the approach similar to that described in our previous paper (Manyukhina, Orekhova, Prokofyev, Obukhova, & Stroganova, 2022). Stimuli and the experimental procedure are shown in Figure 1. Stimuli consisted of drifting vertical sinusoidal gratings (1 cycle/degree, 4 degrees/sec) covered by two-dimensional Gaussian envelope with the radius defining the stimulus size (small: 1°, medium: 2.5°, or large: 12°). Participants sat at 60 cm distance from the monitor (Acer XF250Q, 24,5′′ W LED, 1,920× 1,080 resolution, 240 Hz). Stimuli were presented at full contrast. A direction of motion (left or right) was determined randomly for each trial. Participants were instructed to make a two-alternative forced-choice response, indicating the perceived direction of motion (left or right) by pressing the corresponding arrow key on the keyboard, with no time constraints. The inter-trial interval was 500 ms. In the beginning of each trial, a central dot flickered at the screen (50 ms on, 50 ms off, 250 ms on, 150 ms off) followed by the stimulus presentation. The initial stimulus duration was set to 200 ms. The duration was further adjusted depending on a participant’s response using three (one for a stimulus of each size) interleaved one-up two down staircases that converged on 71% correct performance. The duration changed in discrete time steps. The step was initially 12.3 ms and changed to 8.2 ms after the first two reversals and further to 4.1 ms after two more reversals. The session continued until all staircases completed 9 reversals. Each subject had two sessions separated by a short break. In three participants (all with VSS) the data from the second session were lost due to a technical failure.

**Figure 1.**
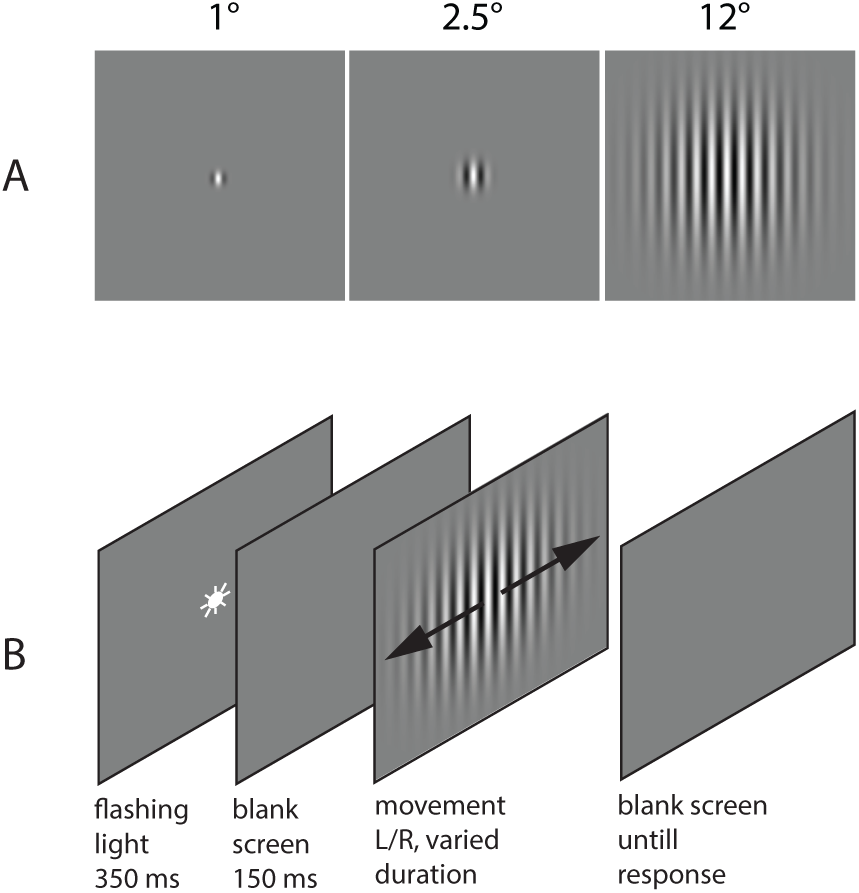
Psychophysical experiment: stimuli (A) and schematic representation of the experimental paradigm. Figure is adapted from (Orekhova et al., 2020), Copyright Elsevier (2020).

For each stimulus size/staircase the threshold was computed by averaging over the reversals, excluding the first two reversals. The reversals followed by 12 or more correct responses were considered as not reflecting the true limit of perceptual capacity, because the probability of accidentally giving 12 correct responses in a row is very low (p=0.00024). These reversals were also excluded from averaging. If there were less than 3 reversals left to calculate the threshold, the staircase was considered a non-converging staircase, and the threshold for the corresponding staircase was not calculated. The spatial suppression index was calculated based on the thresholds for the large and small gratings as follows:

*SSI* = log10(Threshold_Large_) – log10(Threshold_Small_). Larger SSI values indicate stronger perceptual spatial suppression.

### Statistical Analysis

To assess reproducibility of duration thresholds between the two sessions we calculated intraclass correlations using ‘irr’ package in R (two-way mixed effects, absolute agreement). A mixed effects ANOVA was used to assess differences between groups. Duration thresholds were log-transformed to normalize the distributions. Spearman correlation coefficients were calculated to investigate the relationships between psychometric variables and visual perception scores, with Bonferroni correction applied for multiple comparisons to adjust the p-values. The alpha level was set at p < 0.05.

## Results

### Visual symptoms in VSS patients

Participants with VSS had a number of additional visual symptoms. Symptoms critical for the diagnosis of VSS (Puledda et al., 2018) are listed in Table 2. Apart from VS, all of our VSS participants had high prominence of at least one of these critical symptoms (4 or 5 on the Likert scale) and all of them had some additional visual symptoms (e.g. flashes of lights, swirls or clouds with eyes closed, etc.). Besides, patients with VSS had higher scores on the Visual Discomfort Scale than the control participants (t(46) = 6.2, p<0.000001).

### Estimation of duration thresholds

The staircases failed to converge in 5 subjects in the first session and in 2 subjects in the second session, resulting in 2 non-converging staircases for the large grating, 5 non-converging staircases for the medium-sized grating and 2 non-converging staircases for the small gratings. In 7 of 9 cases, the staircases failed to converge during the first session. If the staircase failed to converge in one of the sessions, the threshold was defined based on the results from another session, which was the case in seven subjects, all with VSS.

### Illusion of reverse motion

In 3 of 23 participants with VSS (13%) and in 3 of 28 control participants (11%), the large moving grating elicited a sustained illusion of reverse motion. This illusion was evident in both sessions and precluded assessment of duration thresholds for large-sized gratings. The original psychometric staircases for these participants are presented in the *Supplementary Results*. The illusion tended to disappear at long presentation times (∼1 sec or more). None of the subjects demonstrated the reverse motion illusion in the case of the medium- or small-sized gratings.

### Reliability of duration thresholds assessment

To estimate the reliability of duration thresholds we calculated intraclass correlations (ICC) of the log-transformed duration thresholds between the two sessions. As we were interested in the reliability of the duration thresholds across sessions, we used a model assuming that both the subject and the sessions are random effects (Koo & Li, 2016). In both groups, reliability was good for large gratings and moderate for small gratings, while it was moderate to poor for the medium-sized gratings (Table 3, Figure 2). This result is consistent with the results obtained in our previous study using the same stimuli in an independent sample of healthy adult participants (Orekhova et al., 2020).

**Table 3.** Between-sessions infraclass correlations# of duration thresholds and 95% confidence intervals.

|  | *Large* | *Medium* | *Small* |
| --- | --- | --- | --- |
| VSS | $R_{n=16} = 0.83^{***} (0.59-0.94)$ | $R_{n=17} = 0.56^* (0.14-0.82)$ | $R_{n=16} = 0.61^* (0.2-0.84)$ |
| Control | $R_{n=24} = 0.82^{***} (0.55-0.93)$ | $R_{n=27} = 0.33^* (-0.02-0.62)$ | $R_{n=27} = 0.73^{***} (0.49-0.87)$ |
| VSS + Control | $R_{n=40} = 0.83^{***} (0.64-0.91)$ | $R_{n=44} = 0.48^* (0.22-0.68)$ | $R_{n=43} = 0.69^{***} (0.49-0.83)$ |

<sup>#</sup> Two-way random effects, agreement. \* $p < 0.05$ , \*\*\* $p < 0.0001$
The number of subjects included in the analysis depends on the availability of data from two sessions.

**Figure 2.**
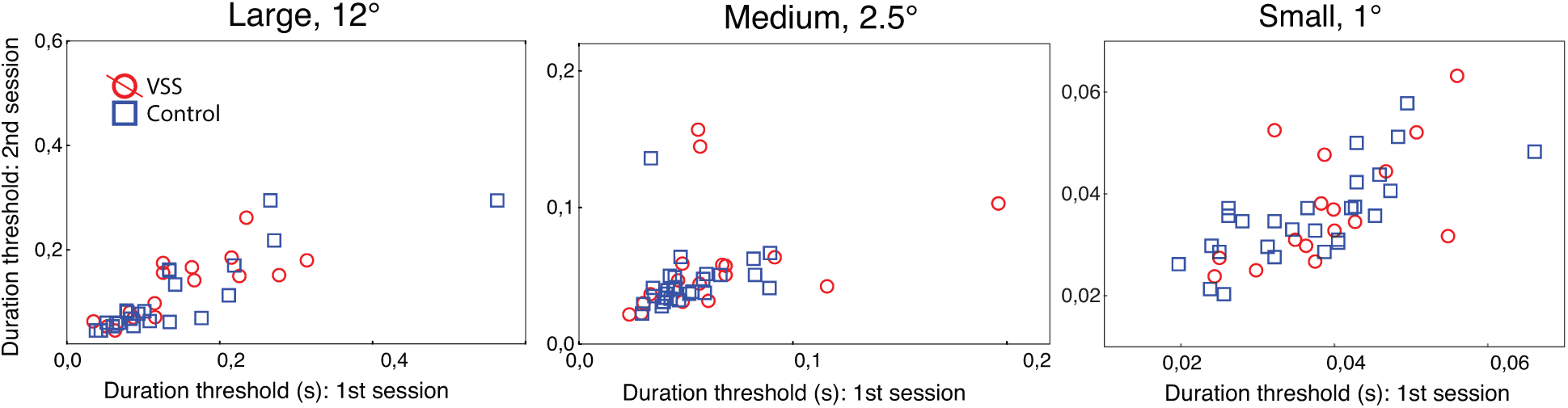
Scatterplots showing between-session consistency of the duration thresholds in VSS and control participants.

### Effect of session order

Mixed ANOVA with the factors Size, Group and Session revealed a significant Session effect (F(1,33)=6.0, p=0.02, partial eta-squared = 0.15) due to an improvement (decrease) in thresholds in the second session compared to the first, probably as a result of practice. The interactions of Session with other factors were not significant (all p>0.4).

When available, the individual thresholds were averaged across sessions for further analysis. However, considering the improvement of performance with practice, we presented the results of group comparison and correlations with visual-perceptual scores for the second session separately in the *Supplementary Results*.

### Group comparison

The medians and range values for duration thresholds in VSS and control participants are summarized in Table 4. Figure 3 shows individual log-transformed duration thresholds. One participant with VSS had an extremely high duration threshold for the small grating (239 ms). This subject (male, 32 years old) was excluded from the ANOVA (but not other analyses). After his exclusion, Levene’s test for homogeneity of variances did not reveal group differences for duration thresholds (Large: F(1,42) = 0.08, p = 0.78; Medium: F(1,48) = 3.05, p = 0.09; Small: F(1,47) = 0.07, p = 0.79).

**Table 4.** Duration thresholds in VSS and control participants: median values and ranges.

| *Grating size* | *VSS* | *Control* |
| --- | --- | --- |
| Large (12°)* | n = 20, M = 100 ms (48 – 248 ms) | n = 24, M = 88 ms (42 – 429 ms) |
| Medium (2.5°) | n = 23, M = 50 ms (23 – 149 ms) | n = 27, M = 44 ms (26 – 88 ms) |
| Small (1°) | n = 23, M = 39 ms (19 – 239 ms) | n = 27, M = 35 ms (23 – 57 ms) |

\* Duration thresholds for the large stimulus were not calculated in participants with the reverse motion illusion.

**Figure 3.**
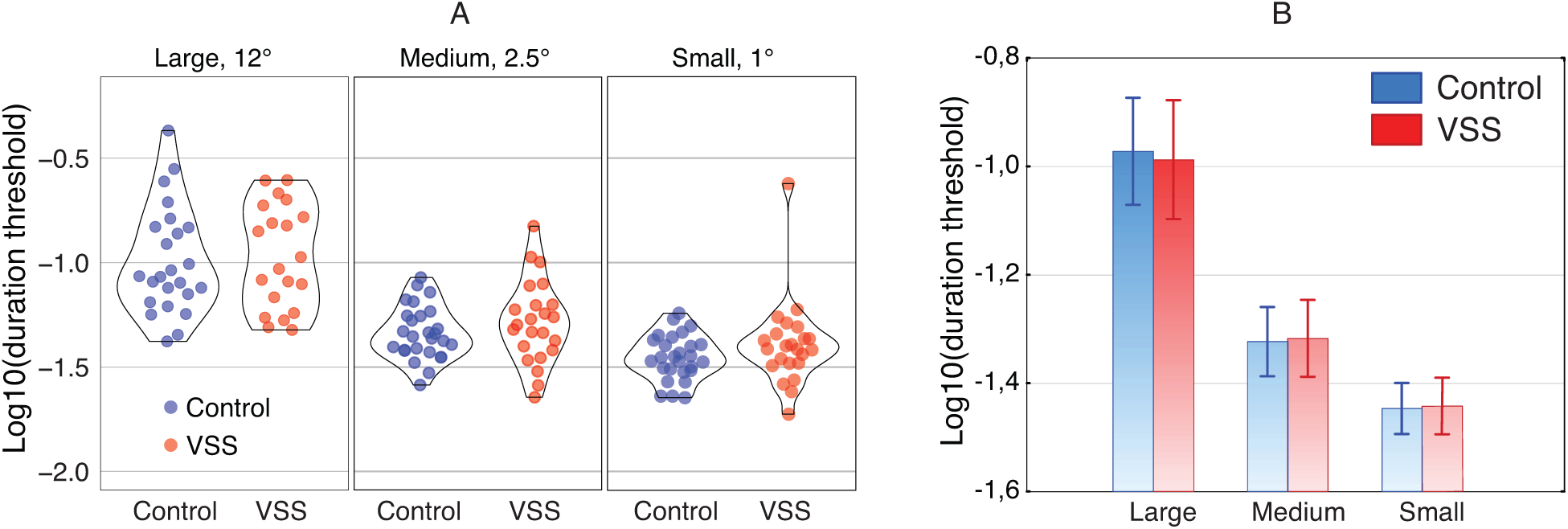
Duration thresholds in VSS and control participants. (A) Between-subject variability in VSS and control groups. (B) Mean duration thresholds and 95% confidence intervals for the subjects included in ANOVA.

To estimate putative group differences in duration thresholds and spatial suppression we performed mixed effects ANOVA with Size as within-unit factor. The between-unit factors were Group, Gender and Group x Gender interaction. We included Gender, since it has been previously shown that males have generally lower duration thresholds than females (Murray et al., 2018).

There was a strong effect of Size (F(2,78) = 120.9, epsilon = 0.80, p<0.00001; partial eta-squared = 0.76), but no effect of Group (F(1,39) = 0.001, p=0.97; partial eta-squared = 0.00) or Size x Group (F(2,78) = 0.08, epsilon = 0.80, p=0.89; partial eta-squared = 0.00) interaction, indicating that the VSS and control groups, on average, had similar duration thresholds and exhibited equally strong spatial suppression. There was a significant main effect of Gender (F(1,39) = 8.8, p = 0.005; partial eta-squared = 0.18): the duration thresholds were shorter in males. Interactions of Gender with other factors were not significant (all p > 0.15). Thus, the ANOVA results indicated no significant differences between the VSS and control groups in either duration thresholds or the Spatial Suppression Index (SSI). A previous study demonstrated that surround suppression is increased in individuals experiencing migraine, including those with and without aura (Battista, Badcock, & McKendrick, 2011). The majority of patients with VSS (17 out of 23) had migraines. However, the SSI in these 17 VSS patients did not differ from that in control participants (F(1,37) = 0.34, p = 0.57, partial eta-squared = 0.01).

### Correlations between duration thresholds and visual disturbances in people with VSS

Table 5 presents Spearman correlations between visual symptoms and performance on the psychometric task. There was a strong correlation between scores on the Palinopsia-II scale (TTP) and duration thresholds for small and medium-size gratings (small: rho = 0.72, uncorrected p = 0.0004; medium: rho = 0.63, uncorrected p = 0.0012; both correlations survive Bonferroni correction for multiple comparisons). As illustrated in Figure 4, VSS subjects with TTP had lower duration thresholds compared to those VSS subjects who experienced less or no TTP symptoms.

**Table 5.**
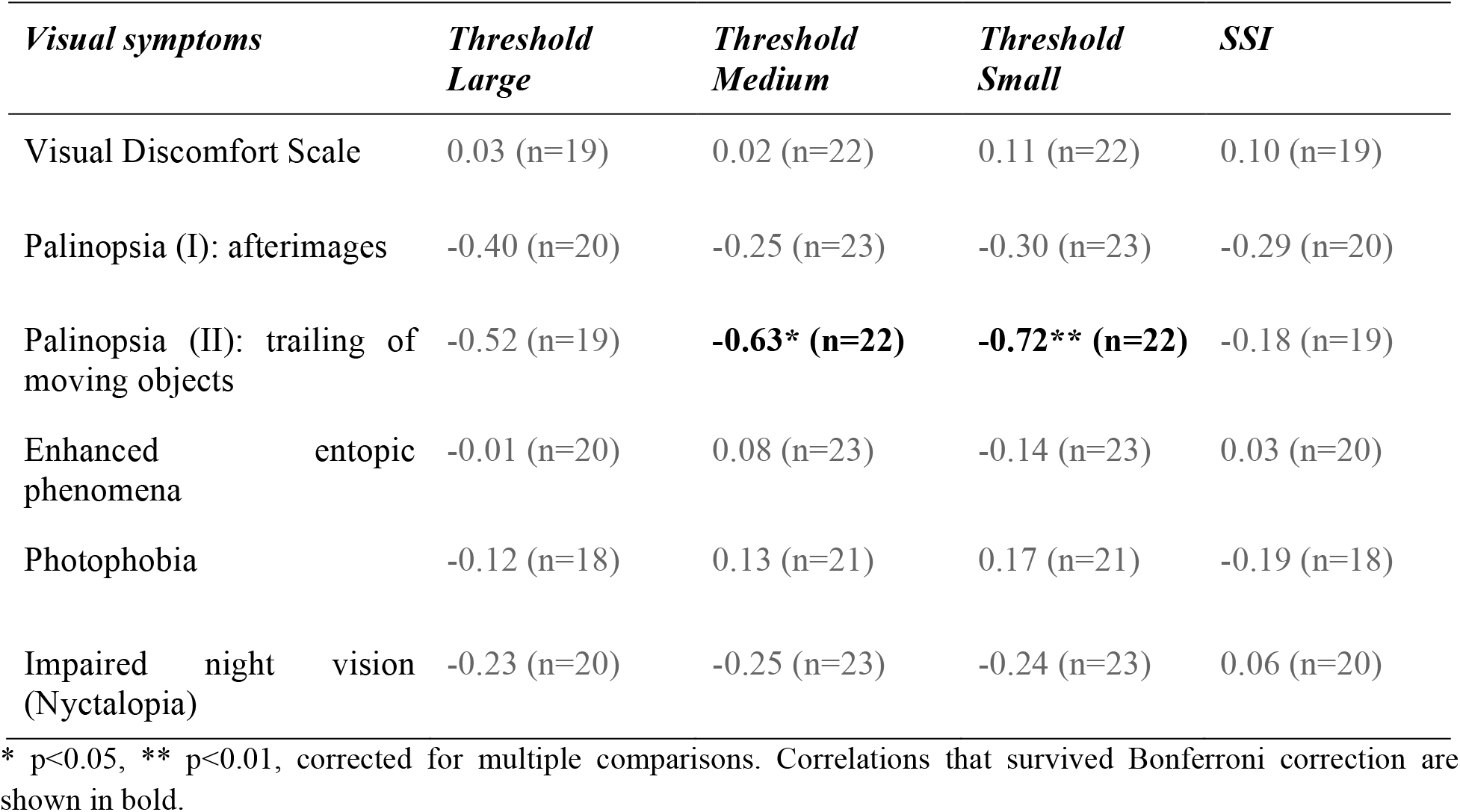
Spearman correlation between the severity of visual disturbances and visual task performance in patients with VSS.

**Figure 4.**
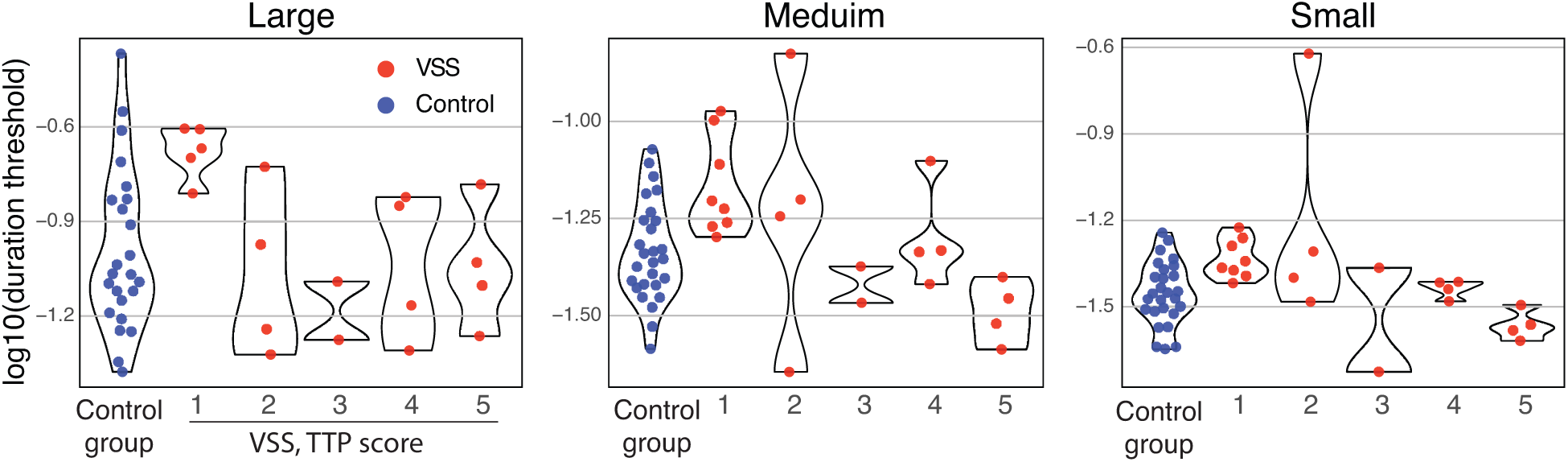
Duration thresholds for the gratings of different sizes in control participants and in participants with VSS, split by scores on palinopsia-II (trailing type) scale. The higher scores denote the greater severity of trailing-type palinopsia

When compared to control participants, VSS patients without trailing-type palinopsia (scored ‘1’ on the Likert scale) exhibited significantly elevated duration thresholds for all stimulus sizes (Small: n = 27/8, Z = -2.7, p = 0.006; Medium: n = 27/8, Z = -2.8, p = 0.006; Large: n = 24/5, Z = -2.6, p = 0.01). On the other hand, in those VSS subjects who scored 4 or 5 on the TTP scale the duration thresholds did not differ from normal (Small: n = 27/8, Z = 1.04, p = 0.29; Medium: n = 27/8, Z = 1.10, p = 0.27; Large: n = 24/8, Z = 0.28, p = 0.77).

Notably, although duration thresholds correlated with TTP scores in VSS patients, they remained within the range of variation observed in control subjects. Specifically, participants without TTP had thresholds overlapping with the higher end of the normal range, while those with TTP aligned with the lower end. The only exception was a VSS patient without TTP (scored “2” on the trailing-type palinopsia scale), who had an extremely high perceptual duration threshold for small-size gratings (Figure 3).

Given the significant effect of gender on duration thresholds (with shorter thresholds in males), we examined the male-to-female ratio in the groups of subjects who scored ‘5’ and ‘1’ on the TTP scale.. Among the patients who scored ‘5’, there were two males and two females. Among the patients who scored ‘1’, there were three males and five females. We also calculated correlations with TTP scores separately for males and females. All correlations were negative and, in the case of small gratings, significant in both males and females (p<0.05, uncorrected for multiple comparisons, see *Supplementary Results*).

## Discussion

A putative imbalance between excitation and inhibition in visual cortical areas may alter visual motion processing in patients with VSS, differently affecting the time required to identify a direction of motion of large and small stimuli (i.e., duration thresholds). To investigate this possibility, we measured the duration thresholds for drifting gratings of different sizes in people with VSS and neurologically healthy control participants. Our findings indicate that, as a group, VSS patients do not differ from control participants in either motion duration discrimination thresholds or in their dependence on stimulus size. However, in the VSS group, duration thresholds were significantly influenced by the presence/absence of a palinopsia variant characterized by visual trailing of moving objects.

Animal studies have shown that increasing stimulus size results in activation of the ‘far surround’ of neurons in the primary visual cortex (V1), which in turn leads to suppression of neuronal activity via excitatory feedback from the higher-order cortical areas to V1 inhibitory circuitry (Angelucci et al., 2017; Nurminen, Merlin, Bijanzadeh, Federer, & Angelucci, 2018). The increase in suppressive feedback limiting neuron sensitivity may explain why the time required to discern the direction of motion of a high-contrast iso-oriented patterned stimulus (grating) increases as a function of stimulus size (Tadin et al., 2003). Our participants with VSS exhibited a normal spatial suppression effect, characterized by a worsening of motion duration thresholds with increasing grating size (Figure 3). Additionally, an equal number of individuals in VSS and control groups (3 in each group) exhibited the reverse motion illusion which has been previously reported in case of brief motion of large high-contrast stimuli and was explained by surround suppression effects (Glasser, Tadin, & Pack, 2014). In sum, these findings suggest that VS symptoms are not necessarily associated with abnormal center-surround suppression caused by the motion of large stimuli.

This result contradicts the findings of McKendrick et al. (McKendrick et al., 2017), who observed reduced center-surround contrast suppression in individuals with VSS. The discrepancy between these results could be attributed to differences in experimental paradigms and/or participant variability. Unlike the contrast suppression task used by McKendrick et al., our task may place greater demand on the magnocellular subdivision of the visual pathway, which is more sensitive to motion and may be relatively unaffected in VSS. In addition, the small sample size in McKendrick’s study (16 VSS and 18 control participants) cannot entirely rule out the possibility of sampling bias, which can occasionally skew the distribution of individual measures of surround suppression. Future studies with larger and more diverse samples are needed to clarify these differences.

Although 74% of our VSS participants had migraine, their typical spatial suppression distinguished them from migraine patients without VSS. Battista and colleagues found heightened spatial suppression in individuals with migraine, which they attributed to an enhanced top-down feedback signaling (Battista et al., 2011). These authors suggested that this enhanced feedback leads to exaggerated surround inhibition, compensating for the hyperexcitability in V1 circuitry. The difference between our results and those of Battista et al suggests that migraine with and without VS may have a different neural basis. While our findings in VSS patients do not rule out potential alterations in excitatory or inhibitory signaling during motion processing, the overall balance between excitatory and inhibitory influences in VSS appears to be preserved, as evidenced by their normal duration thresholds and normal suppressive effect of increasing stimulus size.

For small stimuli that do not extend beyond the population receptive field (less than ∼ 1 degree of visual angle for the central visual field; (Dumoulin & Wandell, 2008)), directional sensitivity is primarily determined by local excitatory and inhibitory processes (Angelucci et al., 2017). Thus, a direct interpretation of the fact that the perceptual duration thresholds for small gratings (Figure 3) are normal in VSS is that the VS illusion, a symptom that was characteristic of all individuals with VSS, is not due to a general inhibitory deficit in the early visual cortex. This interpretation is consistent with the finding of normal luminance increment thresholds for small stationary gratings in patients with VSS in a luminance detection task (Brooks et al., 2022). Indeed, reduced lateral inhibition within the classical receptive fields in V1 is known to decrease discrimination accuracy for moving stimuli (Nelson, Toth, Sheth, & Sur, 1994) and alters contrast sensitivity (Cone, Scantlen, Histed, & Maunsell, 2019; Zhou, Shi, Peng, Hua, & Hua, 2011). The presence of normal duration thresholds for small gratings in our VSS participants is incompatible with the “neural noise” hypothesis, which posits that VSS arises from overall increased spontaneous activity in V1 (Metzler & Robertson, 2018), as the excessive noise would impair direction sensitivity to the small moving stimuli (Lee et al., 2012; Shadlen & Newsome, 1994).

Although duration thresholds in VSS patients were normal at the group level, they correlated negatively with the severity of comorbid TTP symptoms (Table 5). Specifically, greater TTP severity was associated with relatively shorter (improved) duration thresholds, suggesting that higher TTP symptom intensity may enhance directional sensitivity. Of note, none of the VSS patients who demonstrated the illusion of reverse motion during the presentation of large gratings had TTP: all three of them scored ‘1’ on the 5-points Likert scale. Given the lack of prior studies using spatial suppression paradigm in individuals with TTP, either in conjunction with or independent of VSS, our findings regarding the link between TTP and motion duration thresholds, though preliminary, are particularly intriguing and merit further investigation. The neural mechanisms underlying the positive aftereffect of motion in TTP remain largely unknown. Regardless of the specific mechanism that preserves perceptual representations after a moving object has disappeared, this phenomenon may contribute to reduced motion duration thresholds for drifting gratings by effectively extending the perceived exposure time. In this way, trailing-type palinopsia might counterbalance the relatively worsened duration thresholds observed in our participants with VSS without TTP, potentially mitigating the effects of local E/I imbalances. However, it is important to note that the small sample size in our study warrants replication in future research. Moving forward, the incorporation of direction discrimination tasks with varying contrast or noise levels but relatively long exposure times (e.g., (Rudolph & Pasternak, 1999)) could provide valuable insights. Paired with the exposure duration-dependent thresholds that we used in the current study, this combined approach will allow for a more comprehensive assessment of visual motion sensitivity in patients with VSS and palinopsia, furthering our understanding of their altered motion perception.

## Conclusions

In conclusion, our findings suggest that the VS illusion, the hallmark symptom of VSS, is not associated with impaired motion direction discrimination or abnormal center-surround suppressive interactions modulated by input from the higher-tier cortical areas. Although VS frequently co-occurs with palinopsia, the presence vs absence of trailing-type palinopsia seems to exert a qualitatively distinct influence on motion discrimination sensitivity in VSS patients. The variability introduced by this comorbid condition emphasizes the need to account for individual differences in this comorbidity in neurophysiological and clinical models of VSS, as the presence of TTP may significantly affect experimental outcomes.

## Supporting information

Supplementary Results

