## Supplementary Results for "Directional motion sensitivity in people with Visual Snow Syndrome is modulated by the presence of trailing-type palinopsia"

#### 1. Duration thresholds in 3 VSS and 3 control participants experiencing illusion of reverse motion during viewing motion of the large (12°) gratings: original staircases.

Control subject #1, session1

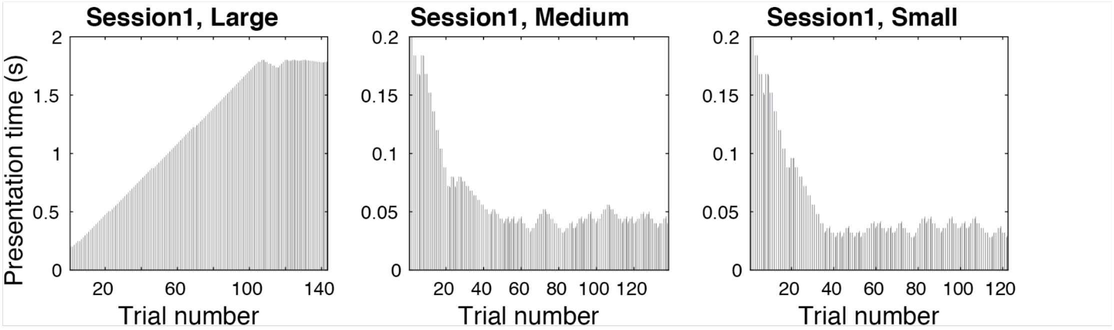


Control subject #1, session2

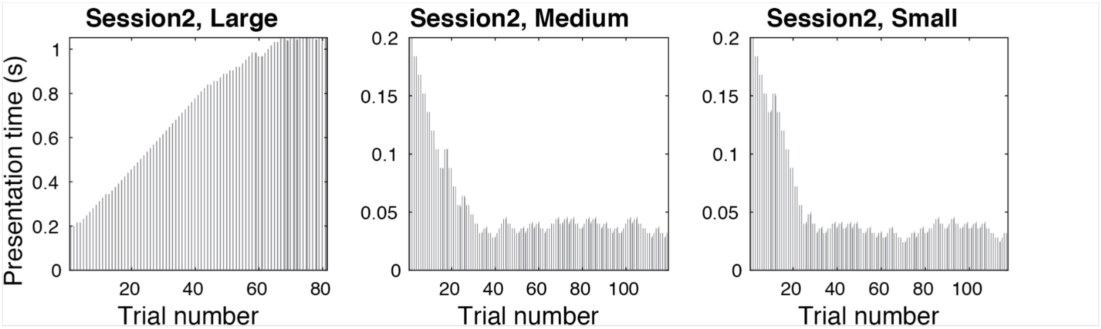

Control subject #2, session 1

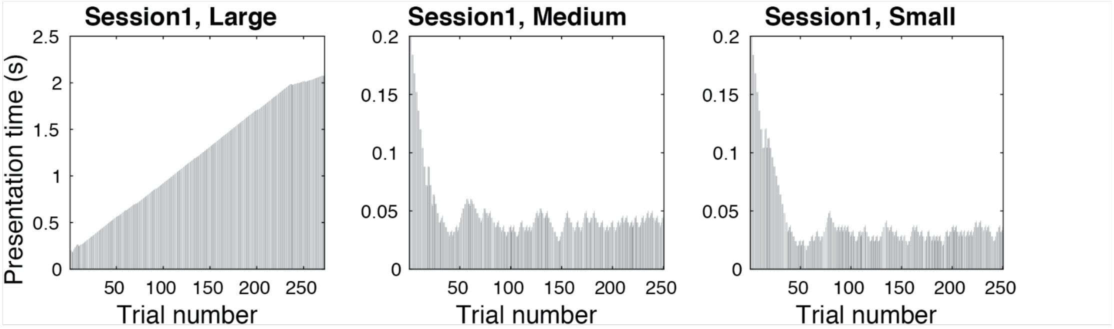


Control subject #2, session 2

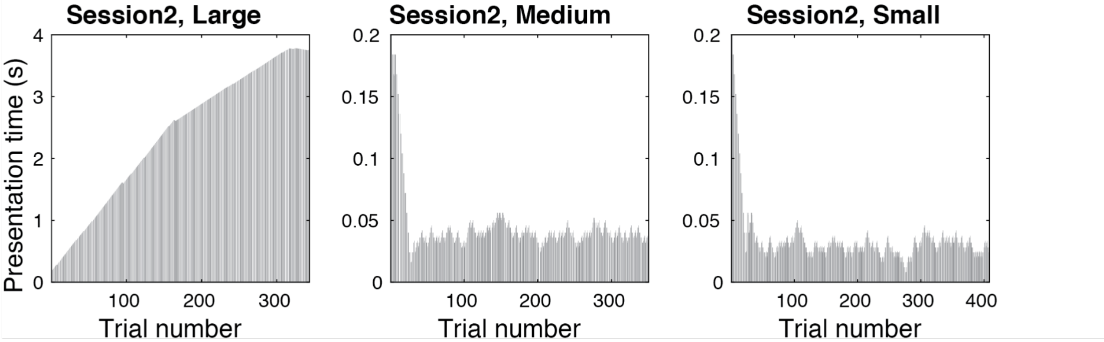


Control subject#3, session 1

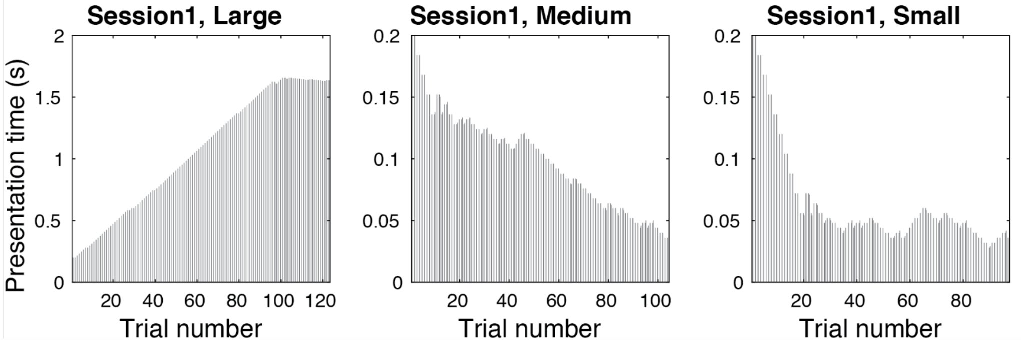


Control subject#3, session 2

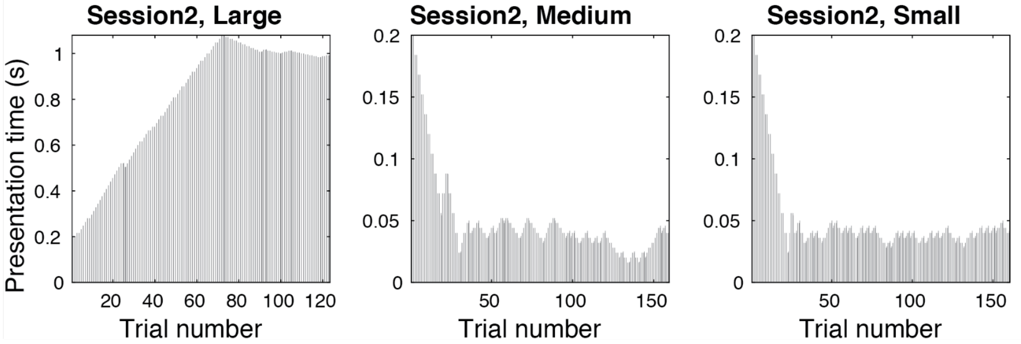

### VSS subject #1, session 1

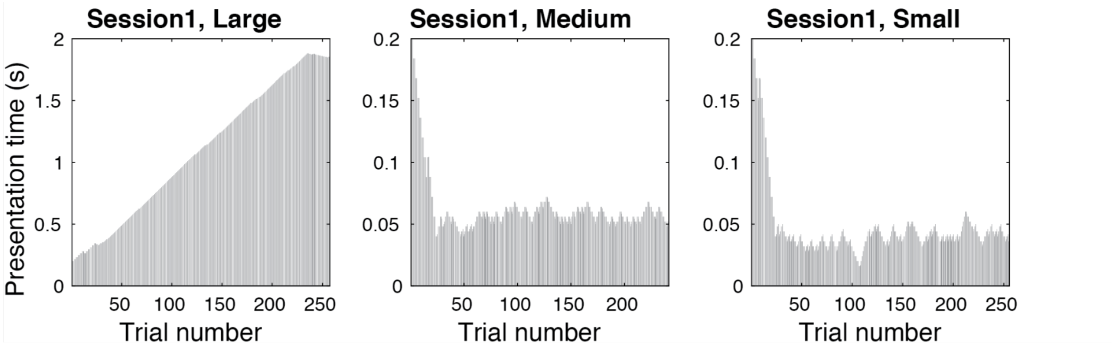


### VSS subject #1, session 2

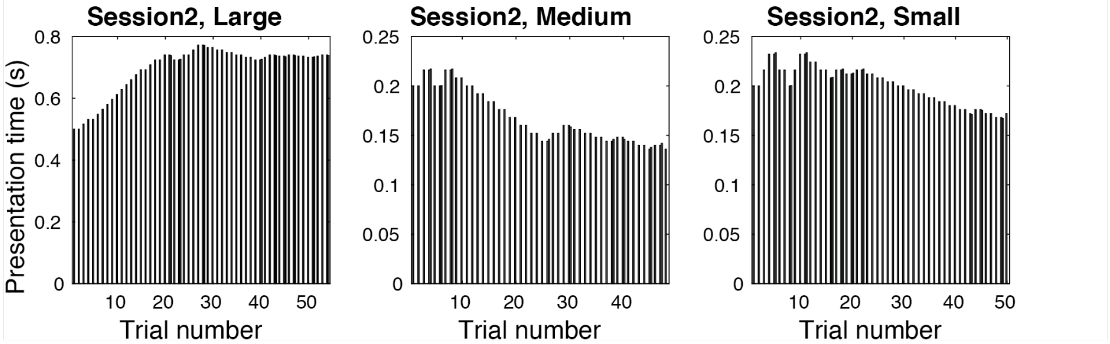


Note: For the #1 VSS participant, in the 2nd session, the starting presentation time for the Large grating was 400 ms.

### VSS subject #2, session 1

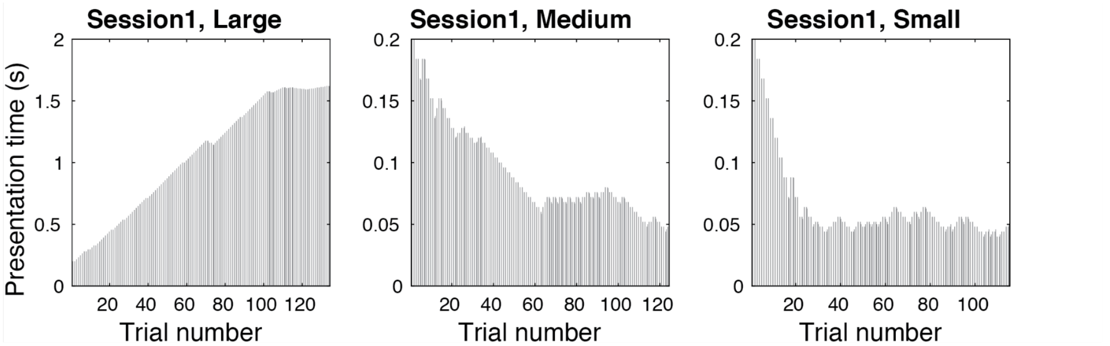


### VSS subject #2, session 2

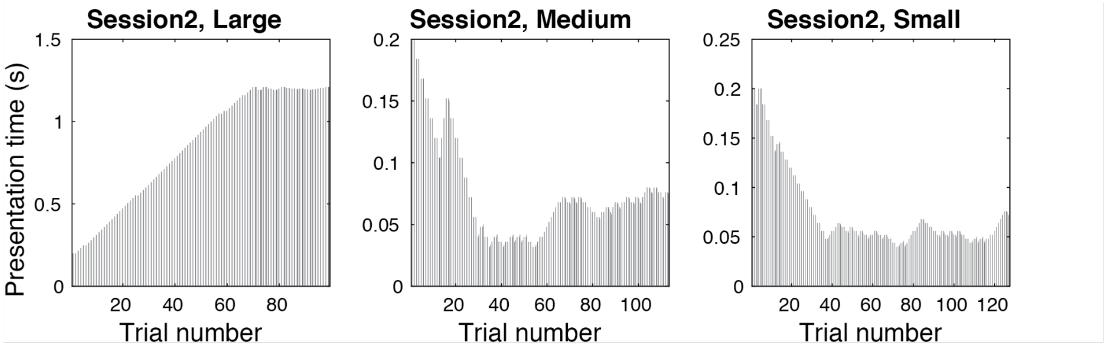

VSS subject #3, session 1

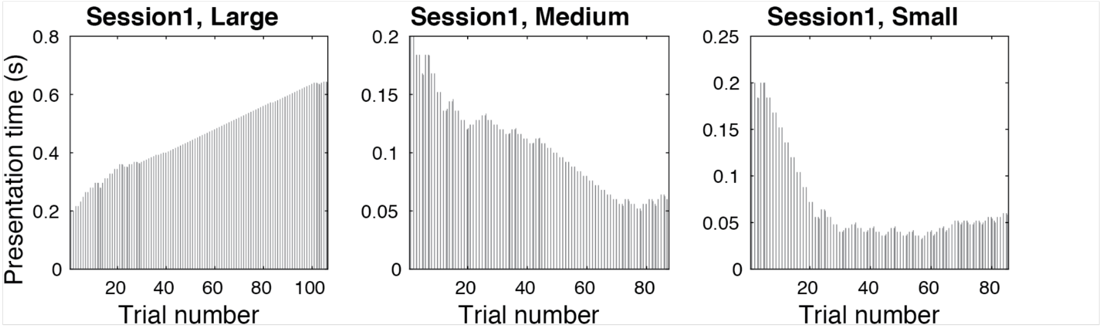


VSS subject #3, session 2

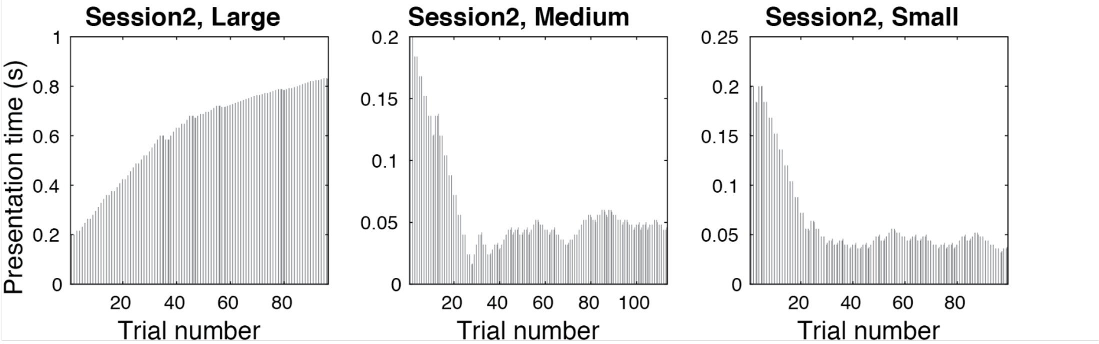

### 2. Results based on the thresholds estimated during the second session

*Group comparison*

The analysis repeated for the second session (in 16 VSS and 22 control subjects) produced the results similar to those for the full data (effect of Size: F(2,72) = 110.3, epsilon = 0.95, p<0.00001; partial eta-squared = 0.75; effect of Gender: F(1,36) = 8.4, p = 0.006; partial eta-squared = 0.19; no significant effects including Group).

*Correlations between psychometric variables and visual disturbances in people with VSS*

Correlations with the trailing-type palinopsia scale were also observed for thresholds assessed during second session (i.e., after training). A positive correlation between strength of spatial suppression estimated by SSI and scores on the visual discomfort scale did not survive correction for multiple comparisons.

**Table 1S.** Spearman correlations between the severity of visual disturbances and visual task performance in patients with VSS in the second session.

| Visual symptoms | Threshold Large | Threshold Medium | Threshold Small | SSI |
|---|---|---|---|---|
| Visual Discomfort Scale | 0.45 (n=16) | 0.13 (n=18) | 0.1 (N=16) | 0.54 (n=16) |
| Palinopsia (I): afterimages | -0.27 (n=18) | -0.47 (n=20) | -0.51 (n=18) | -0.10 (n=18) |
| Palinopsia (II): trailing of moving objects | -0.59 (n=18) | **-0.76* (n=20)** | **-0.82* (n=18)** | -0.33 (n=18) |
| Enhanced entopic phenomena | 0.20 (n=18) | -0.08 (n=20) | -0.25 (n=18) | 0.32 (n=18) |
| Photophobia | -0.01 (n=17) | 0.11 (n=19) | 0.13 (n=17) | -.10 (n=17) |
| Impaired night vision (Nyctalopia) | 0.03 (n-18) | -0.29 (n=20) | -0.14 (n=18) | 0.15 (n=18) |

* p<0.05, corrected for multiple comparisons. Correlations that survived Bonferroni corrections are shown in bold.

### 3. Correlations between duration thresholds and severity of trailing-type palinopsia (TTP) in male and female VSS patients.

**Table S2.** Spearman correlations between TTP scores and duration thresholds in patients with VSS.

| *Visual symptoms* | *Threshold Large* | *Threshold Medium* | *Threshold Small* |
|---|---|---|---|
| Males | -0.20 (n=10) | -0.47 (n=12) | 0.76** (n=12) |
| Females | -0.84** (n=9) | -0.83** (n=10) | 0.64* (n=10) |

* p<0.05, ** p<0.01, uncorrected for multiple comparisons.
